# [Re] Optimization of a free water elimination two-compartment model for diffusion tensor imaging

**DOI:** 10.1101/108795

**Authors:** Rafael Neto Henriques, Ariel Rokem, Eleftherios Garyfallidis, Samuel St-Jean, Eric Thomas Peterson, Marta Morgado Correia

**Affiliations:** MRC Cognition and Brain Sciences Unit, Cambridge, Cambridgeshire, UK; The University of Washington eScience Institute, Seattle, WA, USA; Intelligent Systems Engineering, Indiana University, IA, USA; University Medical Center Utrecht, Utrecht, NL; Biosciences, SRI International, Menlo Park, CA, USA

## Abstract

Typical diffusion-weighted imaging (DWI) is susceptible to partial volume effects: different types of tissue that reside in the same voxel are
inextricably mixed. For instance, in regions near the cerebral ventricles or parenchyma, fractional anisotropy (FA) from diffusion tensor imaging (DTI) may be underestimated, due to partial volumes of cerebral spinal fluid (CSF). Free-water can be suppressed by adding parameters to diffusion MRI models. For example, the DTI model can be extended to separately take into account the contributions of tissue and CSF, by representing the tissue compartment with an anisotropic diffusion tensor and the CSF compartment as an isotropic free water diffusion coefficient. Recently, two procedures were proposed to fit this two-compartment model to diffusion-weighted data acquired for at least two different non-zero diffusion MRI b-values. In this work, the first open-source reference implementation of these procedures is provided. In addition to presenting some methodological improvements that increase model fitting robustness, the free water DTI procedures are re-evaluated using Monte-Carlo multicompartmental simulations. Analogous to previous studies, our results show that the free water elimination DTI model is able to remove confounding effects of fast diffusion for typical FA values of brain white matter. In addition, this study confirms that for a fixed scanning time the fwDTI fitting procedures have better performance when data is acquired for diffusion gradient direction evenly distributed along two b-values of 500 and 1500 s/mm^2^.

## Introduction

Diffusion-weighted Magnetic Resonance Imaging (DW-MRI) is a biomedical imaging technique that allows for the non-invasive acquisition of *in vivo* data from which tissue microstructure can be inferred. Diffusion tensor imaging (DTI), one of the most commonly used DW-MRI techniques in the brain, models diffusion anisotropy of tissues using a second-order tensor known as the diffusion tensor (DT) [1], [2].

DTI-based measures such as fractional anisotropy (FA) and mean diffusivity (MD) are usually used to assess properties of brain microstructure. For example, FA is sensitive to different microstructural properties (such as packing density of axons, and density of myelin in nerve fibers [3]), as well to the degree of white matter coherence (i.e. the alignment of axons within a measurement voxel). However, because a measurement voxel can contain partial volumes of different types of tissue, these measures are not always specific to one particular type of tissue. In particular, diffusion anisotropy in voxels near cerebral ventricles and parenchyma can be underestimated by partial volume effects of cerebrospinal fluid (CSF).

Since CSF partial volume effects depend on the exact positioning of brain relative to the MRI spatial encoding grid, the reproducibility of diffusion-weighted measures might be compromised by free water partial volume effects [8]. Moreover, free water contaminations are likely to confound diffusion-based measures in studies of brain aging and some pathologies, because CSF partial volume effects might be increased due to tissue morphological atrophy and enlargement of cerebral ventricle [9].

During the MRI acquisition, the signal from free water can be directly eliminated using fluid-attenuated inversion recovery diffusion-weighted sequences [10]. However, these sequences have some drawbacks. For example, they require longer acquisition times and produce images with lower signal-to-noise ratio (SNR). Moreover, these approaches cannot be used retrospectively for data cohorts already acquired with multi b-value diffusion-weighted sequences, such as the Human Connectome Project [15] and the Cambridge Centre for Ageing and Neuroscience cohort [14].

Free-water can also be suppressed by adding parameters to diffusion MRI models. For example, the DTI model can be extended to separately take into account the contributions of tissue and CSF, by representing the tissue compartment with an anisotropic diffusion tensor and the CSF compartment as an isotropic free water diffusion coefficient [11]. In addition to the correction for free water partial volume effects, estimates of volume fraction provided for this model were shown to provide insight into the brain as an indicator of extracellular space which changes due to concussions [12]. Recently, two procedures were proposed by Hoy and colleagues to fit this two-compartment model to diffusion-weighted data acquired for at least two different non-zero b-values (i.e. different diffusion gradient-weightings or diffusion shells) [5]. Although these procedures have been shown to provide diffusion-based measures that are stable to different degrees of free water contamination, the authors noted that their original algorithms were “implemented by a group member with no formal programming training and without optimization for speed” [5].

In this work, we provide the first open-source reference implementation of the free water contamination DTI model (see function.py). All implementations are made in Python, and are designed to be compatible with previously optimized functions that are freely available as part of the open-source software package Diffusion Imaging in Python (Dipy, [4]). In addition to presenting some methodological improvements that increase model fitting robustness, we also present here an evaluation of the fitting that replicates the results of Hoy et al. (2014)[5]. Finally, the optimal acquisition parameter sets for fitting the free water elimination DTI model are re-assessed.

## Methods

The free water elimination DTI (fwDTI) model describes the measured diffusion-weighted signal *s*_*i*_ with a simple bi-exponential expansion of DTI:

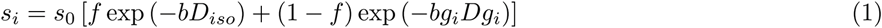

where *s*_0_ is the signal when no diffusion sensitization gradient is applied, *f* is the volume fraction of free water contamination, *D* is the diffusion tensor of the tissue, *b* is a parameter that quantifies the amount of diffusion sensitization, *g*_*i*_ is the diffusion gradient direction and *D*_*iso*_ is the free water isotropic diffusion coefficient which is set to 3.0 × 10^−3^*mm*^2^/*s* [11].

### Implementation of fitting algorithms

Since no source implementation was previously provided by Hoy and colleagues, our implementation relies on the equations provided in the original article.

#### Weighted-Linear Least Square (WLS)

Two corrections to the equations of the first algorithm are proposed. Firstly, the free-water adjusted diffusion-weighted signal formula (original article’s Methods subsection “FWE-DTI”) should be written as:

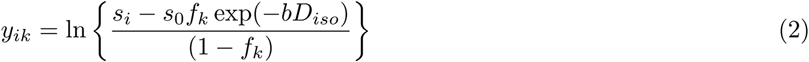

instead of:

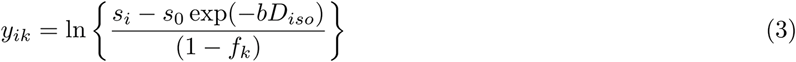

where *f*_*k*_ is a grid search free water volume fraction sample.

Secondly, according to the general linear least squares solution [6], the parameters matrix is estimated using the weighted linear least squares solution of the free-water elimination model:

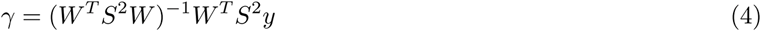

instead of:

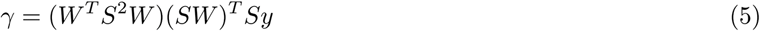

where *γ* contains the fwDTI model parameters 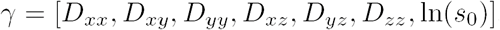, *y* is a matrix containing the elements of *y*_*ik*_ computed from equation 2, *S* is a diagonal matrix with diagonal set to the *s*_*i*_ samples, and *W* is a matrix computed from the *m* diffusion-weighted directions *g*_*i*_ and b-values:

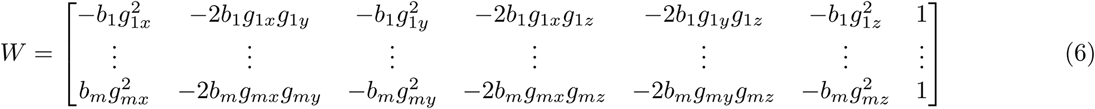

Relative to the original article, the elements of *W* have been re-ordered according to the order of the diffusion tensor elements used in Dipy. This reordering is just a matter of differing notational conventions and has no effect on the results of the equation.

To ensure that the WLS method converges to the local minimum, *f* grid search sampling is performed over larger interval ranges relative to the original article. Particularly, for the second and third iterations used to refine the parameters precision, *f* is resampled over intervals of 0.2 and 0.02 instead of interval sizes of 0.1 and 0.01 proposed by Hoy and colleagues. On the other hand, the sample step size was maintained to 0.01 and 0.001 respectively.

Moreover, since the WLS objective function is sensitive to the squared error of the model weights (*ω*_*i*_ = *s*_*i*_):

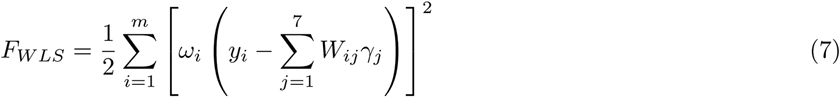

when evaluating which (*f*, *D*_*tissue*_) pair is associated with smaller residuals the NLS objective function is used instead:

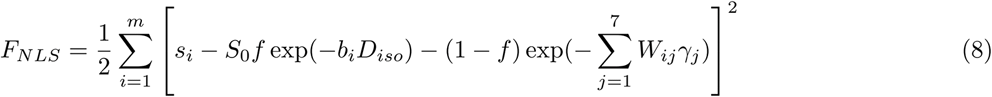

Similarly to the original article [5], this procedure is only used to obtain the initial guess for the free water elimination parameters, which were then used to initialize a fwDTI model non-linear convergence solver (see below).

#### Removing problematic first guess estimates

For cases where the ground truth free water volume fraction is close to one (i.e. voxels containing only free water), the tissue’s diffusion tensor component can erroneously fit the free water diffusion signal and therefore the free water DTI model fails to identify the signal as pure free water. In these cases, since both tissue and free water compartments can represent the signal well, the free water volume fraction can be arbitrarily estimated to any value in the range between 0 and 1 rather than being close to 1. To overcome this issue, before the use of the non-linear fitting procedure, Hoy et al. (2014) proposed to re-initialize the parameters initial guess when the tissue’s mean diffusivity approaches the value of free water diffusion (tissue’s MD > 1.5 × 10^−3^*mm*^2^/*s*) [5]. In the present study, these re-initialization settings are re-evaluated by assessing the fwDTI estimates for synthetic voxels that mostly contain free water diffusion. The frequency with which this problem arises is also analyzed as a function of ground truth *f* values (supplementary_notebook_1.ipynb). Based on this evaluation, the following parameter adjustments were done in our implementations when the initial tissue’s mean diffusivity is higher than 1.5 × 10^−3^
*mm*^2^/*s*: 1) free water volume fraction estimates are set to 1 indicating that the voxel signal is likely to be only related to free water diffusion; 2) the elements of the tissues diffusion tensor is set zero to avoid high tissue’s mean diffusivity for high volume fraction estimates and to better indicate that tissue diffusion is practically absent.

#### Non-Linear Least Square Solution (NLS)

To improve computation speed, non-linear convergence was implemented using Scipy’s wrapped modified Levenberg-Marquardt algorithm (function `scipy.optimize.leastsq` of Scipy). This replaces the use of a modified Newton’s algorithm proposed in the original article,

To constrain the model parameters to plausible ranges, some variable transformations can be applied to the nonlinear objective function. These were implemented as optional features that can be controlled through user-provided arguments. To restrict the range of the volume fraction to values between 0 and 1, the variable *f* in equation 8 can be replaced by sin(*f*_*t*_ – *π*/2)/2 + 1/2 and non-linear convergence is then performed as a function of *f*_*t*_. To ensure that the diffusion tensor is positive definite, diffusion parameters can be converted to the Cholesky decomposition elements as described in [7].

In addition to the `scipy.optimize.leastsq` function, a more recently implemented version of Scipy’s optimization function `scipy.optimize.least_square` (available as of Scipy’s version 0.17) was also tested. The latter directly solves the non-linear problem with predefined constraints in a similar fashion to what is done in the original article. However, our experiments showed that this procedure does not exceed the performance of `scipy.optimize.leastsq` in terms of accuracy, and requires more computing time (see supplementary_notebook_2.ipynb for more details).

To speed up the performance of the non-linear optimization procedure, the Jacobian of the free water elimination DTI model was analytically derived and incorporated in the non-linear procedure (for details of the Jacobian derivation see supplementary_notebook_3.ipynb). Due to increased mathematical complexity, our analytical Jacobian derivation is not compatible with the Cholesky decomposition. This variable transformation is therefore not used by default.

#### Implementation dependencies

In addition to the dependency on Scipy, both free water elimination fitting procedures require modules from Dipy [4], which contain implementations of standard diffusion tensor fitting functions. An installation of Dipy is therefore required, before the fwDTI fitting procedures proposed here can be used. To install Dipy please follow the steps described in dipy’s website, or follow the instructions in the README file of the code provided with this paper. In addition to Dipy’s modules, the implemented procedures also require the NumPy Python package, which is also a dependency of both Scipy and Dipy.

### Simulation 1

The performance of the fwDTI fitting techniques was first assessed for five different tissues’ diffusion tensor FA levels and for free water volume fraction contamination using a Monte Carlo simulation of a multi-tensor implemented in Dipy. This simulation corresponds to the results reported in Figure 5 of the original article [5].

The acquisition parameters for this first simulation were based on the previously optimized parameter set reported by Hoy et al. (2014) which consist in a total of 70 signal measured for diffusion b-values of 500 and 1500 *s*/*mm*^2^ (each along 32 different diffusion directions) and for six b-value=0 *s*/*mm*^2^. As in the original article, the ground truth of tissue’s diffusion tensor had a constant diffusion trace of 2.4 × 10^−3^*mm*^2^/*s*. The diffusion tensor’s eigenvalues used for the five FA levels are reported in Table 1.

**Table 1.** Eigenvalues values used for the simulations

| FA | 0 | 0.11 | 0.22 | 0.3 | 0.71 |
| --- | --- | --- | --- | --- | --- |
| $\lambda_1$ | $8.00 \times 10^{-4}$ | $9.00 \times 10^{-4}$ | $1.00 \times 10^{-3}$ | $1.08 \times 10^{-3}$ | $1.60 \times 10^{-3}$ |
| $\lambda_2$ | $8.00 \times 10^{-4}$ | $7.63 \times 10^{-4}$ | $7.25 \times 10^{-4}$ | $6.95 \times 10^{-4}$ | $5.00 \times 10^{-4}$ |
| $\lambda_3$ | $8.00 \times 10^{-4}$ | $7.38 \times 10^{-4}$ | $6.75 \times 10^{-4}$ | $6.25 \times 10^{-4}$ | $3.00 \times 10^{-4}$ |

For each FA value, eleven different degrees of free water contamination were evaluated (f values equally spaced from 0 to 1). To assess the procedure’s robustness to MRI noise, Rician noise with signal-to-noise ratio (SNR) of 40 relative to the b-value=0 *s*/*mm*^2^ images was used, similarly to Hoy et al. (2014) [5]. For each FA and f-value pair, simulations were performed for 120 different diffusion tensor orientations which were repeated 100 times for a total of 12000 simulated iterations for each FA and f-value pair [5].

### Simulation 2

After evaluating the fitting procedures for the reported FA and *f* ground truth values, the optimal b-values for two-shell diffusion MRI acquisition were re-assessed. This second simulation corresponds to the original article’s Figure 2 [5] in which b-value minimum was investigated from 200 to 800 *s*/*mm*^2^ in increments of 100 *s*/*mm*^2^ while b-value maximum was assessed from 300 to 1500 *s*/*mm*^2^ in increments of 100 *s*/*mm*^2^. Similarly to Hoy et al. (2014), this simulation was done for a fixed f-value of 0.5 and FA of 0.71, corresponding to a typical MRI voxel in the boundary between CSF and white matter tissue. For each b-value pair, simulations were repeated 12000 times similarly to simulation 1 and fitting performances was quantified from the free water DTI’s FA estimates, *f* estimates, and mean diffusivity estimates, using the mean squared error: 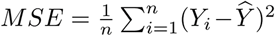, where *n*=12000, *Ŷ* is the ground truth value of a diffusion measure, and *Y*_*i*
_ is single measure estimate.

### Simulation 3

To evaluate our implementations for different number of diffusion shells, different number of gradient directions, and different SNR levels, the lower panels of the original article’s Figure 4 were replicated. For this last simulation, multi-tensor simulations were performed for six different acquisition parameters sets that are summarized in Table 2.

**Table 2:** The six different acquisition parameter set tested

| Number of shells | b-values ($s/mm^2$) | Directions per shell |
| --- | --- | --- |
| 2 | 500/1500 | 32 per shell |
| 3 | 500/1000/1500 | 21/21/22 |
| 4 | 400/767/1133/1500 | 16 per shell |
| 6 | 400/620/840/1060/1280/1500 | 10/10/10/10/10/14 |
| 8 | 300/471/643/814/986/1157/1329/1500 | 8 per shell |
| 16 | 300/380/460/540/620/700/780/860/940/<br>1020/1100/1180/1250/1340/1420/1500 | 4 per shell |

Although Hoy et al. (2014) only reported the latter analysis for the higher FA level, here the lower FA values of 0 was also assessed. According to the original article, this optimization analysis was also done for the fixed f-value of 0.5. As the previous two simulations, tests were repeated 12000 times (for 120 diffusion tensor direction each repeated 100 times). Fitting performances was quantified using the mean squared error of the the free water DTI FA, *f* and MD estimates (see Simulation 2).

### *In vivo* data

Similarly to the original article, the procedures were also tested using *in vivo* human brain data [13], that can be automatically downloaded using functions implemented in Dipy functions (see code script). The original dataset consisted of 74 volumes of images acquired for a b-value of 0 *s*/*mm*^2^ and 578 volumes diffusion weighted images acquired along 16 diffusion gradient directions for b-values of 200 and 400 *s*/*mm*^2^ and along 182 diffusion gradient directions for b-values of 1000, 2000 and 3000 *s*/*mm*^2^. In this study, only the data for b-values up to 2000 *s*/*mm*^2^ were used, to decrease the impact of non-Gaussian diffusion more apparent at higher b-values. This is because these effects are not included in the free water elimination model [5]. We also processed the data with the standard DTI tensor model (as implemented in Dipy), in order to compare the results with the free water elimination model.

## Results

The results from the first multi-tensor simulations are shown in Figure 1. As reported in the original article, no FA bias was observed for large FA ground truth values and free water volume fractions *f* ranging around 0 to 0.7 (top panel of Figure 1). However, FA values seemed to be overestimated for higher volume fractions. This bias was more prominent for lower FA values in which overestimations wee visible starting from lower free water volume fractions. The results shown in the lower panels of Figure 1 suggest that the free water elimination model accurately estimated the free water volume fraction for the full range of volume fraction ground truth values. All the features observed here are consistent with Figure 5 of the original article.

**Figure 1:**
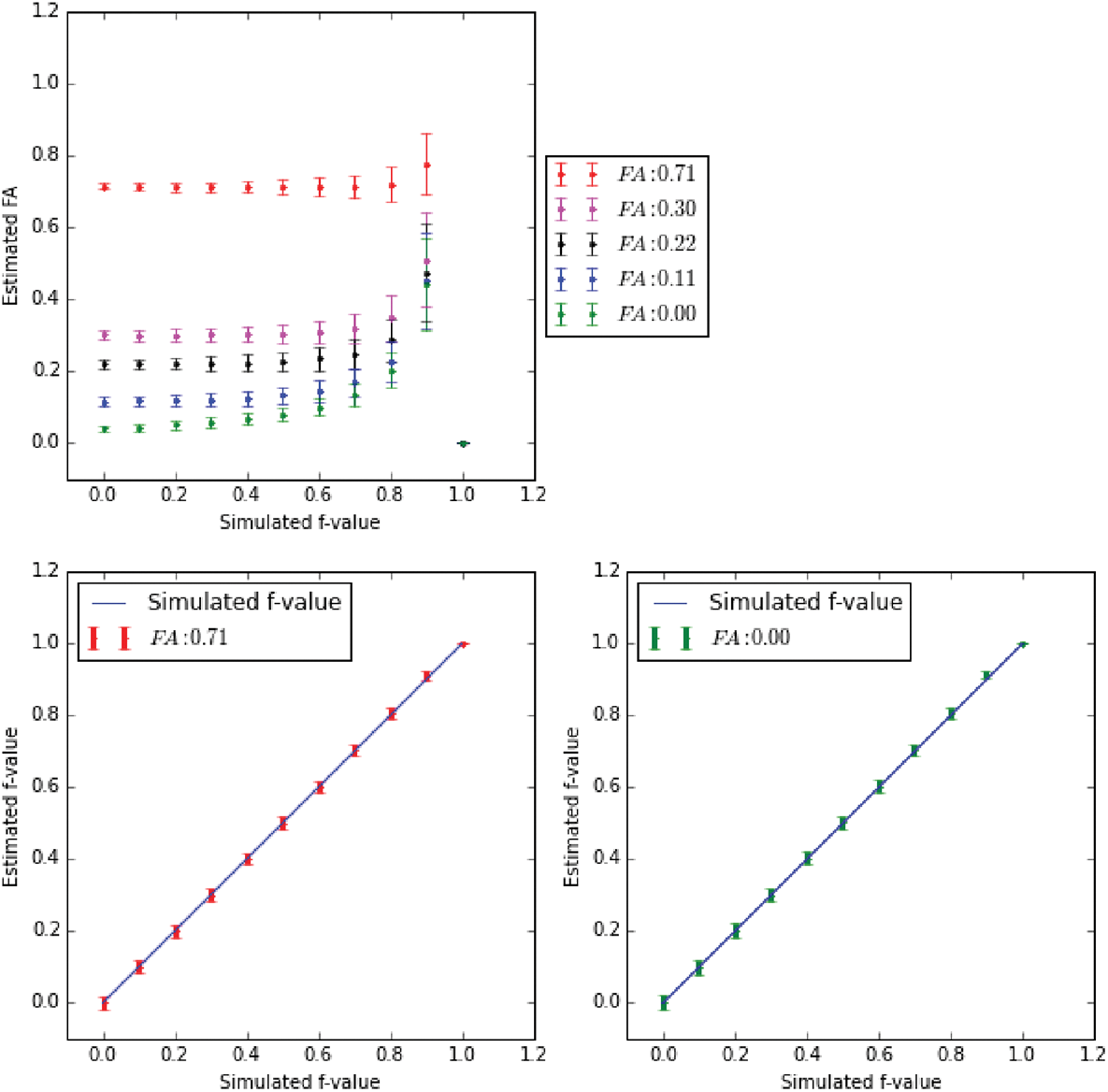
Fractional Anisotropy (FA) and free water volume fraction (*f*) estimates obtained with multi-tensor simulations using the free water elimination fitting procedures. The top panel shows the FA median and interquartile range for the five different FA ground truth levels and plotted as a function of the ground truth water volume fraction. The bottom panels show the estimated volume fraction *f* median and interquartile range as a function of its ground truth values (right and left panels correspond to the higher and lower FA values, respectively). This figure reproduces Fig. 5 of the original article.

The re-evaluation of the optimal b-value for a two-shell acquisition scheme is shown in Figure 2. For a better visualization of the optimal b-value pair, MSE is converted to it reciprocal inverse [5]. In this way, the optimal b-value pair corresponds to a inverse reciprocal mean squared error of 1 (IRMSE). Although our fwDTI fitting implementations seem to produce more stable IRMSE for all diffusion measures (note that colormaps where rescaled from 0.9 to 1), results of Figure 2 are consistent to the original article, suggesting that the b-values pair 500-1500 *s*/*mm*^2^ is the most adequate parameter setting for fwDTI model fitting.

**Figure 2:**
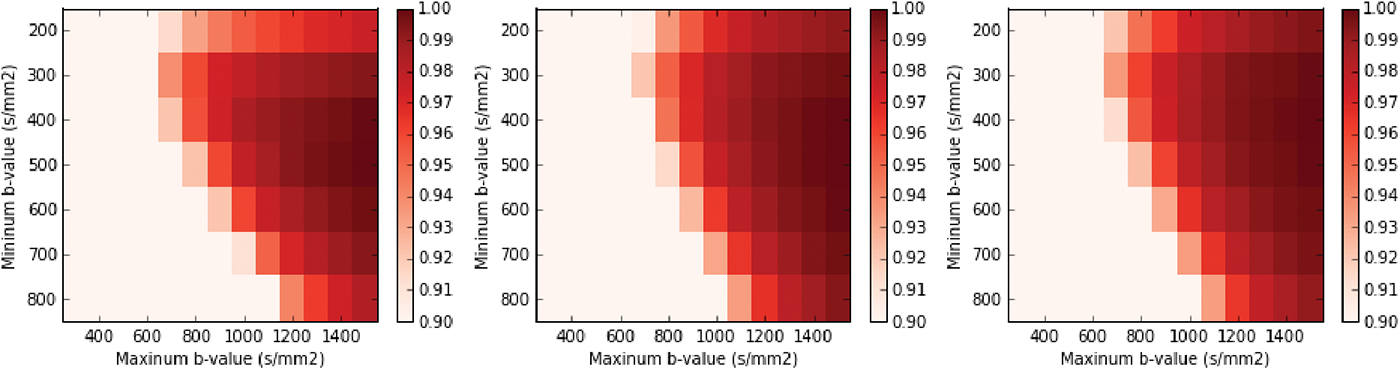
Inverse reciprocal mean squared errors (IRMSE) as a function of the minimum and maximum b-value for three different diffusion measures. For left to right panels, the IRMSE were computed from the tissue’s fractional anisotropy, free water volume fraction estimates, and tissue’s mean diffusivity. MD colormaps are in ×10^−3^*mm*^2^/*s*. This figure reproduces Fig. 2 of the original article.

Figure 3 shows the mean squared errors as a function of the signal to noise ratio (SNR) for different sets of acquisition parameters: different number of b-value shells and gradient directions. As reported in the original article [5], upper panels show that the two-shell acquisition scheme provides better diffusion measure estimates, when the ground truth FA is high. This suggests that it is preferable to use a small number of different b-values (shells) with a large number of directions per b-value, rather than using a large number of b-values with a smaller number of directions in each b-value shell. With the exception of the MSE for FA estimates, results are consistent for the ground truth FA=0 (lower panels of Figure 3). The similar performances of fwDTI in estimating tissue FA for the different parameter sets is expected. This is because in this case tissue has isotropic diffusion, FA estimates should not depend on the number of sampled directions.

**Figure 3:**
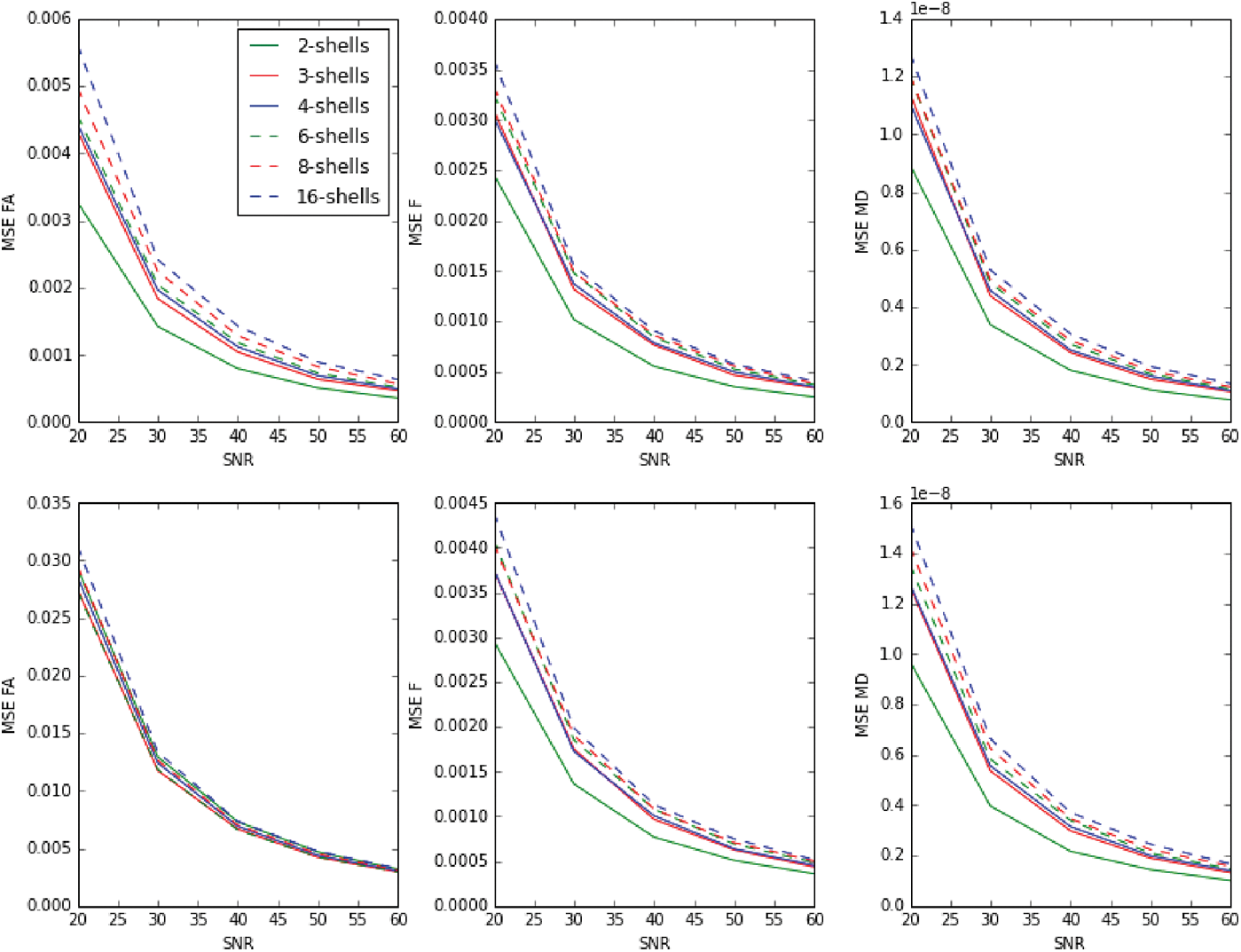
Mean squared errors (MSE) plotted as a function of the signal to noise ratio (SNR) for six different acquisition parameter sets. For left to right panels the MSE were computed from the tissue fractional anisotropy estimates, free water volume fraction estimates and tissue mean diffusivity. The upper panels reproduce the lower panels of the original article Fig. 4, in which ground truth tissue FA is set to 0.71. Lower panels show the analogous results for a ground truth tissue FA of 0

*In vivo* tensor statistics obtained from the free water elimination and standard DTI models are shown in Figure 4. Complete processing of all these measure took less than 1 hour on an average Desktop and Laptop PC (~2GHz processor speed), while the reported processing time by Hoy et al. was around 20 hours. The free water elimination model seems to produce higher values of FA in general and lower values of MD relative to the metrics obtained from the standard DTI model. These differences in FA and MD estimates are expected due to the suppression of the isotropic diffusion of free water. As similarly reported in the original article, high amplitudes of FA are observed in the periventricular gray matter which might be related to inflated values in voxels with high *f* values. These can be mitigated by excluding voxels with high free water volume fraction estimates (see supplementary_notebook_4.ipynb), as suggested by Hoy and colleagues [5].

**Figure 4:**
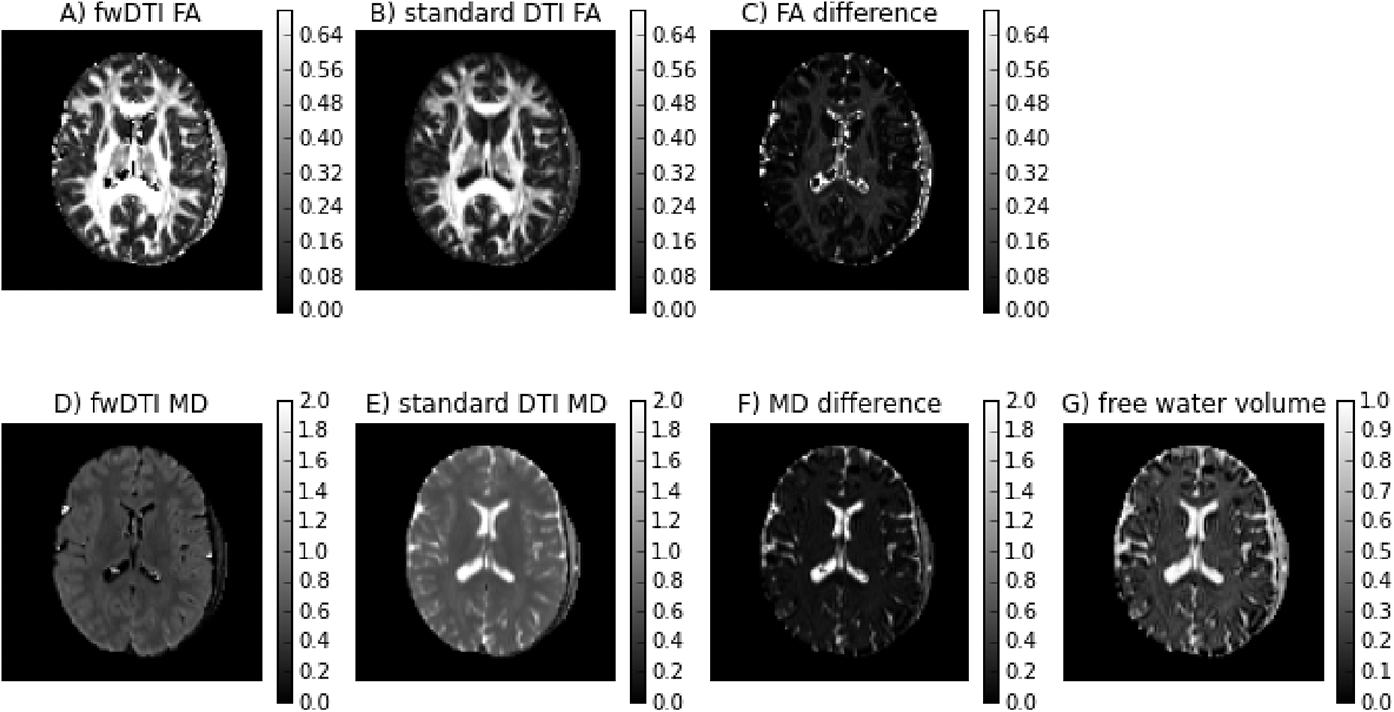
*In vivo* diffusion measures obtained from the free water DTI and standard DTI. The values of FA for the free water DTI model, the standard DTI model and their difference are shown in the top panels (A-C), while respective MD values are shown in the bottom panels (D-F). In addition, the free water volume fraction estimated from the free water DTI model is shown in panel G. This figure reproduces Fig. 7 of the original article.

## Conclusion

Despite the changes done to reduce the algorithm’s execution time, the implemented procedures to solve the free water elimination DTI model have comparable performance in terms of accuracy to the original methods described by Hoy and colleagues [5]. Based on simulations similar to the ones used in the original article, our results confirm that the free water elimination DTI model is able to remove confounding effects of fast diffusion for typical FA values of brain white matter. As previously reported by Hoy and colleagues, the proposed procedures seem to generate biased values of FA for free water volume fractions near one. Nevertheless, our results confirm that these problematic cases correspond to regions of the *in vivo* data that are not typically of interest in neuroimaging analysis (voxels associated with cerebral ventricles) and might be removed by excluding voxels with measured volume fractions above a reasonable threshold such as 0.7. This study also confirms that for a fixed scanning time the fwDTI fitting procedures have better performance when data is acquired for diffusion gradient direction evenly distributed along two b-values of 500 and 1500 *s*/*mm*^2^.

## Author Contributions

Conceptualization: RNH, AR, MMC. Data Curation: RNH, AR, EG, SSTJ. Formal Analysis: RNH. Funding Acquisition: RNH, AR. Investigation: RNH. Methodology: RNH, AR, EG. Project Administration: RNH, MMC, AR, EG. Resources: RNH, MMC, AR. Software: RNH, AR, ETP, EF, SSTJ. Supervision: MMC, AR. Validation: AR, SSTJ, EG. Visualization: RNH. Writing - Original Draft Preparation: RNH. Writing - Review & Editing: RNH, AR, MMC, ETP, SSTJ.

## Acknowledgments

Rafael Neto Henriques was funded by Fundação para a Ciência e Tecnologia FCT/MCE (PIDDAC) under grant SFRH/BD/80114/2012.

Ariel Rokem was funded through a grant from the Gordon & Betty Moore Foundation and the Alfred P. Sloan Foundation to the University of Washington eScience Institute.

Thanks to Romain Valabregue, CENIR, Paris for providing the data used here.

